# Inhibition of GMP synthesis extends yeast replicative lifespan

**DOI:** 10.1101/629428

**Authors:** E. A. Sarnoski, P. Liu, G. Urbonaite, T. T. Olmez, T. Z. Young, R. Song, M. Acar

## Abstract

Aging, the time-dependent accumulation of molecular damage, is the primary limiting factor of human lifespan. Slowing down damage accumulation or its prevention therefore represents a promising therapeutic paradigm to combat aging-related disease and death. While several chemical compounds extend lifespan in model organisms, their mechanism of action is often unknown, reducing their therapeutic potential. Using a systematic approach, here we show that inhibition of GMP synthesis is a novel lifespan extension mechanism in yeast. We further discover that proteasome activation extends lifespan in part through GMP insufficiency. GMP synthesis inhibition exerts its lifespan extension effect independently of the canonical nutrient-sensing and sirtuin pathways regulating lifespan. Exposing longitudinally aging yeast cells to GMP synthesis inhibition in an age-dependent manner, we demonstrate that the lifespan extension by GMP insufficiency is facilitated by slowing, rather than reversing, the aging process in cells. While GMP and its downstream metabolites are involved in many cellular processes in cells, our results rule out the combined effect of global transcription and translation on cellular lifespan. These findings elucidate the involvement of nucleotide metabolism in the aging process. The existence of clinically-approved GMP synthesis inhibitors elicits the potential of a new class of therapeutics for aging-related disorders.

## INTRODUCTION

Aging is the primary risk factor for human morbidity and mortality throughout the developed world^1–3^. Prevention or elimination of the molecular damage that causes aging therefore represents a promising therapeutic paradigm to combat the plethora of diseases that manifest late in life. Target-based drug discovery has advanced several therapeutic candidates to human trials, including metformin^4^ and mTOR inhibitors^5^. While these treatments hold promise, additional interventions are necessary to fully minimize the health effects of advanced age.

In recent years, phenotype-based discovery approaches^6,7^ have identified many compounds capable of extending lifespan in model organisms^1^. These compounds are a starting point for therapeutic development, but the mechanism of action for many of them remains unknown, reducing their therapeutic potential. One such compound is mycophenolic acid (MPA), an FDA-approved immunosuppressant that extends the replicative lifespan (RLS) of the yeast *Saccharomyces cerevisiae*^6^.

Here, we seek to understand the mechanism behind MPA’s ability to extend replicative lifespan. We find that guanine reverses MPA’s pro-longevity effect, indicating that reduction in cellular GMP pools is responsible for the lifespan extension. We then investigate if MPA’s RLS extension mechanism overlaps with previously-described aging pathways, elucidating that it partially overlaps with the effect exerted by activated proteasome. We characterize a set of cellular processes and phenotypes in terms of how they are affected by MPA and whether or not they can facilitate the observed RLS extension.

## RESULTS

### Inhibition of GMP synthesis extends yeast RLS

MPA’s primary, but not exclusive, effect is to reduce cellular GMP pools through inhibition of inosine monophosphate dehydrogenase (IMD), the rate-limiting enzyme in *de novo* GMP synthesis^8,9^ (Fig. 1A). GMP can also be synthesized via a salvage pathway in the presence of exogenous guanine^8,10^.

**Figure 1.**
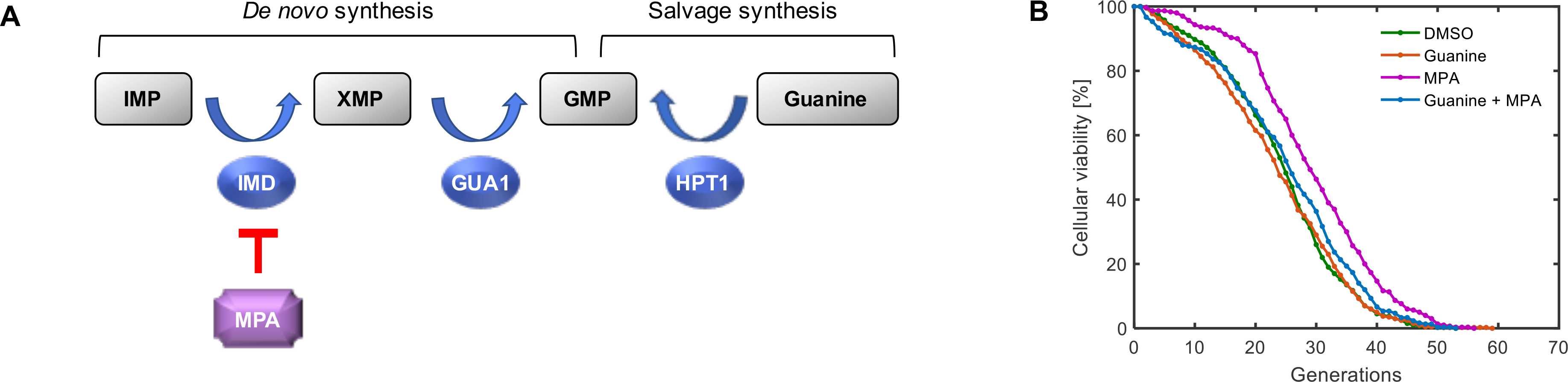
Inhibition of GMP synthesis extends yeast RLS. **A.** Simplified schematic representation of GMP synthesis pathways in *S. cerevisiae*. MPA limits *de novo* GMP synthesis via inhibition of IMD genes. GMP is synthesized via the salvage pathway in the presence of exogenous guanine. **B.** Lifespan curves for wild-type, haploid yeast (BY4741) in the presence or absence of MPA and guanine. N = 400 cells each for the DMSO and guanine conditions, pooled from four independent experiments of 100 cells each; N = 300 cells each for the MPA and guanine&MPA conditions, pooled from three independent experiments of 100 cells each.

As GMP is a nucleotide that is used as a monomer in RNA, we wanted to measure the effect of MPA treatment on transcription by using downstream single-cell gene expression as a probe. For this, we fused GFP to the strongly-expressed *TEF1* promoter and integrated one copy of the P_*TEF1*_-GFP construct in the haploid yeast genome. After growing cells in media containing MPA and/or guanine, we measured the resulting single-cell GFP expression levels (Fig. S3, S4). Adding guanine to the growth media resulted in a 13.7% increase in GFP levels, which we interpret as due to increased mRNA levels because of the role of guanine in mRNA synthesis. On the other hand, growing cells in MPA, and therefore inhibiting GMP synthesis, led to a 15.1% reduction in the GFP expression levels. Finally, adding guanine to the media containing MPA reversed the reduction in GFP levels. To see if these results could be directly attributable to transcription only, we measured total RNA levels in cells grown in the same conditions as above. The resulting global RNA levels supported the trend we saw at the single-cell level using the P_*TEF1*_-GFP probe: adding MPA led to a reduction in the mean RNA levels compared to the DMSO control, and this reduction was reversible by adding guanine to the growth media (Fig. S5, S6).

We hypothesized that the reduction in cellular GMP pools was responsible for MPA’s pro-longevity effect. Using our microfluidic “yeast replicator” platform^11^, we quantified the number of daughter cells produced by 200 mother cells individually tracked throughout their replicative lifespan. These single-cell RLS measurements were made in minimal media environments containing MPA (at standard 10 μM) with or without supplemental guanine. While the average lifespan of the cells grown in the guanine-supplemented environment was slightly reduced compared to DMSO control, the difference was not statistically significant (Table S1). Importantly, we found that the longevity effect of MPA was prevented by the supplementation of exogenous guanine (Fig. 1B). These results confirm that MPA extends replicative lifespan in *S. cerevisiae* through inhibition of GMP synthesis.

### Inhibition of GMP synthesis extends lifespan independent of the nutrient sensing and sirtuin pathways

We next asked how lifespan extension via GMP synthesis inhibition could be placed among the known lifespan pathways. To answer this question, we used a systematic approach to categorize longevity interventions to genetic regulators of lifespan. A longevity intervention can act either within or independent from any known longevity pathway. If within a longevity pathway, the intervention must act upon, upstream from, or downstream of a given pathway component. We aimed to conclusively identify the placement of the lifespan extension mechanism through GMP-synthesis inhibition relative to the known genetic lifespan pathways using the two-step <u>L</u>ongevity <u>P</u>lacement <u>T</u>est (LPT) (Fig. 2A, Fig. S1, Supplementary Information). In Step 1, we ask if the longevity intervention extends lifespan in a strain lacking a critical lifespan pathway component, the probe gene. In Step 2, we ask whether an epistatic agent that prevents lifespan extension from the longevity intervention can also prevent lifespan extension conferred by modulation of the probe gene. By combining this information, one can definitively classify the relationship of a longevity intervention to a known lifespan regulation pathway.

**Figure 2.**
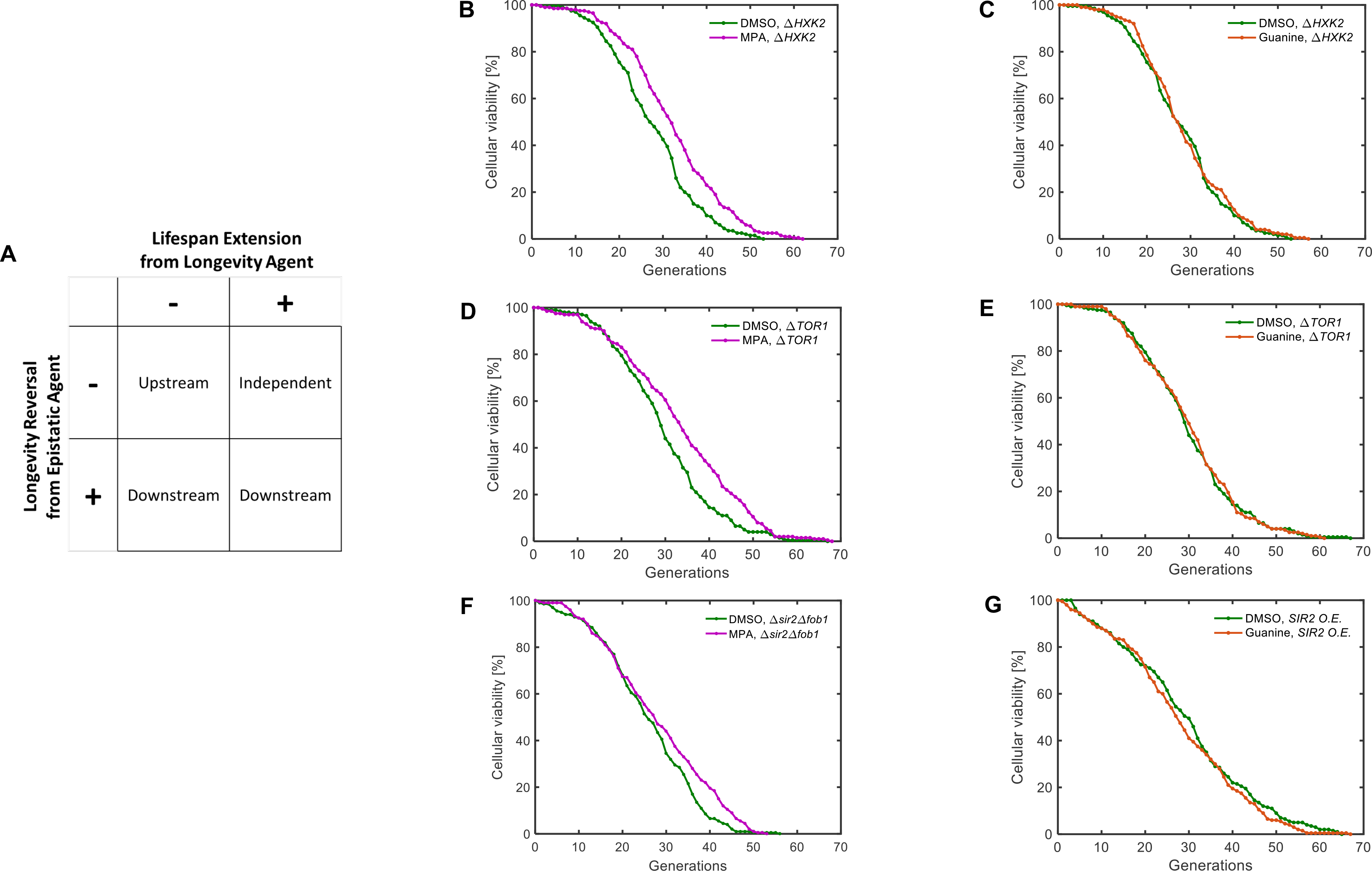
Inhibition of GMP synthesis extends lifespan independent of the nutrient sensing and sirtuin pathways. **A.** Schematic representation of the LPT test and its interpretation. **B-G.** Lifespan curves corresponding to Step 1 (**B**, **D**, **F**) or Step 2 (**C**, **E**, **G**) of the LPT test for the nutrient sensing pathway, including a dietary restriction mimetic (**B**, **C**) and TOR inhibition (**D**, **E**), and the sirtuin pathway (**F**, **G**). N = 200 cells for each condition, pooled from two or more independent experiments.

We used the LPT to determine the relationship of GMP synthesis inhibition to the three major lifespan-extension pathways known for *S. cerevisiae*^12^. For this purpose, MPA was used as the longevity intervention, and guanine was used as the epistatic agent. The first pathway we tested was the nutrient sensing pathway^13^, which encompasses dietary restriction^14^ and target of rapamycin (TOR) inhibition^15^. As the LPT probe genes, we chose TOR1 and HXK2, whose individual deletion is known to extend yeast lifespan. TOR1 is a protein kinase subunit of the TORC1 complex that controls cell growth in response to nutrient availability. HXK2, on the other hand, is a hexokinase whose deletion provides a genetic mimicry of nutrient limitation because it phosphorylates intracellular glucose as part of glucose metabolism. MPA further extended lifespan in the long-lived *ΔTOR1* (*p*=0.000634) and *ΔHXK2* (*p*=0.0000269) strains, while guanine did not suppress gene-deletion-caused lifespan extension in these strains (Fig. 2B-E, Table S1). These results indicate that inhibition of GMP synthesis exerts its longevity effect independent of the nutrient-sensing pathway.

The second lifespan-extension pathway we tested to determine its relationship to the GMP-synthesis inhibition was the sirtuin pathway. As the LPT probe gene, we chose SIR2, an evolutionarily conserved histone deacetylase. SIR2 deletion shortens yeast lifespan, and conversely its overexpression extends lifespan. However, shortened lifespan in the *ΔSIR2* background masks the effect of most longevity interventions because *ΔSIR2* cells die from the rapid accumulation of rDNA circles before other aging factors accumulate^16^. This lifespan shortening and longevity masking can be rescued by concurrent deletion of FOB1^16^, a nucleolar protein whose deletion reduces formation of rDNA circles^17^. MPA was able to extend the lifespan (*p*=0.0123) in the absence of SIR2 in the *ΔSIR2ΔFOB1* background (Fig. 2F) while guanine did not reverse (*p*=0.25) the lifespan extension conferred by SIR2 overexpression (Fig. 2G). These results indicate that inhibition of GMP synthesis extends lifespan independent of the sirtuin pathway.

### Proteasome activation extends lifespan in part through GMP insufficiency

Next, we asked whether inhibition of GMP synthesis extended RLS through the third major lifespan-extension pathway, the proteasome pathway. UBR2 is a ubiquitin ligase subunit, and its deletion activates the proteasome by stabilizing RPN4, a transcription factor that promotes expression of proteasome subunits^18^. Activation of the proteasome via UBR2 deletion extends RLS independent of the nutrient sensing pathway^19^. Therefore, we chose UBR2 as the LPT probe gene. MPA led to a 9.4% extension (*p* = 0.012) of yeast RLS in a *ΔUBR2* strain (Fig. 3A, Table S1), compared to a 22.1% extension (*p* = 1.43*10^−9^) in wild-type cells (Fig. 1B, Table S1). On the other hand, guanine supplementation, at a saturating concentration, partially but significantly (*p* = 0.015) suppressed lifespan extension from UBR2 deletion (Fig. 3B, Table S1). These results indicate that UBR2 deletion activates, if not exclusively, the same lifespan extension mechanism as MPA without saturating the target; in other words, UBR2 deletion extends lifespan in part through GMP insufficiency. A role for GMP metabolism in the phenotypic effects of UBR2 deletion is reinforced by the observation that IMD proteins are among the most highly upregulated proteins in a *ΔUBR2* strain^19^. We therefore conclude that, downstream of UBR2, activated proteasome exerts its lifespan-extension effect partially through GMP insufficiency, as supplementing *ΔUBR2* strain’s growth media with excess guanine significantly but only partially reverses the lifespan extension. These conclusions are further supported by the following observations. While MPA extends the lifespan of wild-type cells by ~5.16 generations and the UBR2 deletion without MPA similarly leads to ~5.4 generations of extension, when we measured the average lifespan of UBR2-deleted cells in the MPA environment, we saw an average lifespan of 31.5 generations (or an 8.11-generation extension compared to wild-type), corresponding to less than the ~10.56-generation extension that would be expected if the effects of MPA and UBR2 deletion were fully non-overlapping.

**Figure 3.**
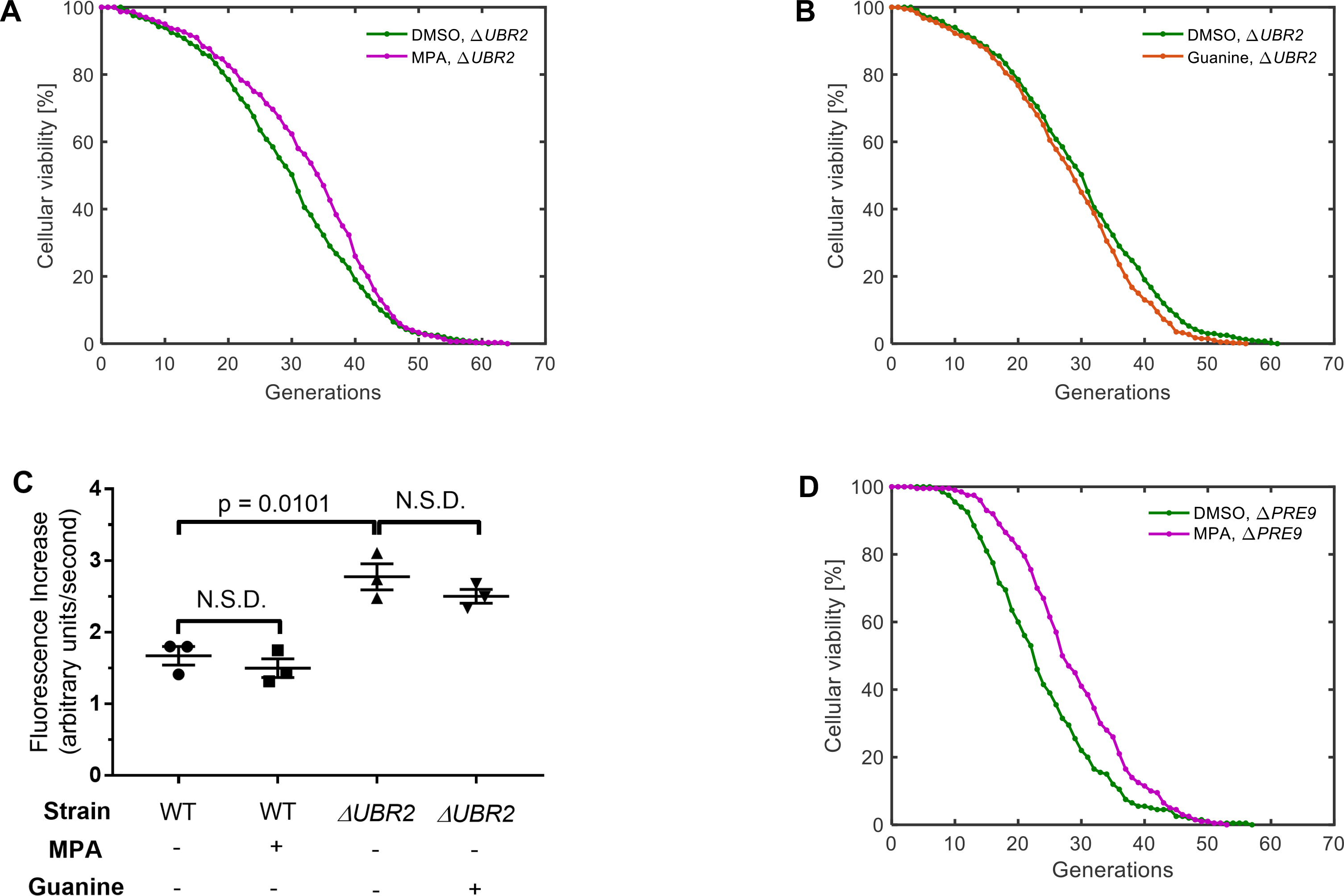
Proteasome activation extends lifespan through GMP inhibition. **A-B.** Lifespan curves corresponding to Step 1 (**A**) and Step 2 (**B**) of the LPT test for the proteasome pathway of lifespan extension. N = 300 cells for the MPA condition, pooled from three independent experiments of 100 cells each; N = 400 cells for the DMSO condition, pooled from four independent experiments of 100 cells each; N = 400 cells for the guanine condition, pooled from four independent experiments of 100 cells each. **C.** Proteasome activity for wild-type (BY4741) cells, or ΔUBR2 cells, in the presence or absence of MPA or guanine. N = 3 biological replicates for each condition. Errors bars are standard error of the mean. NSD, no significant difference. **D.** Lifespan curve for a ΔPRE9 strain in the presence or absence of MPA. N = 200 cells for each condition, pooled from two independent experiments of 100 cells each.

Since multiple steps exist between UBR2 deletion and proteasome activation, it was possible that MPA acted downstream of UBR2, but upstream of proteasome activation. We sought to differentiate these possibilities. We measured proteasome activity in wild-type cells in the presence and absence of MPA (Fig. 3C, Fig. S2), and found that MPA does not activate the proteasome. Furthermore, guanine does not alter proteasome activity in *ΔUBR2* cells (Fig. 3C). This demonstrates that MPA regulates lifespan without modulating the proteasome. We further demonstrated that MPA extends RLS in a *ΔPRE9* strain (Fig. 3D, Table S1), in which proteasome activation is not able to increase RLS due to the absence of the proteasome subunit PRE9^2^. These results indicate that MPA acts to extend lifespan downstream of proteasome activation. Since deletion of UBR2 extends lifespan exclusively^19^ through proteasome activation and excess guanine supplementation only partially reverses the lifespan-extension effect of UBR2 deletion, we conclude that proteasome activation extends lifespan in part through GMP insufficiency. While the other mechanism(s) through which activated proteasome extends lifespan is still unknown, a previous hypothesis^2,3^ suggests that proteasome activation leads to lifespan extension via more efficient removal of misfolded and damaged proteins. Future studies will show which proteins are more prone to aging-associated damage and how their removal through activated proteasome impacts cellular lifespan.

### MPA slows, rather than reverses, the aging process

Interventions that extend lifespan may mechanistically act to slow the accumulation of aging factors or aging-related damage, reverse aging-related damage, or suppress its effects. We sought to determine through which mechanism the inhibition of GMP synthesis extended the lifespan. For this purpose, we assessed the RLS of yeast cells treated with MPA only for the first 24 hours of a Replicator experiment (Fig. 4A), only after the first 24 hours of a Replicator experiment (Fig. 4B), or with a 6-hour pulse treatment between the 24^th^ and 30^th^ hour of the experiment (Fig. 4C). We found that MPA treatment for only part of the lifespan, either early or late, resulted in reduced lifespan extension compared to whole-lifespan treatment (Fig. 4A-B, Table S1). Furthermore, pulse treatment resulted in little to no lifespan extension (Fig. 4C, Table S1). These results indicate that inhibition of GMP synthesis slows, rather than reverses, the accumulation of aging factors in cells.

**Figure 4.**
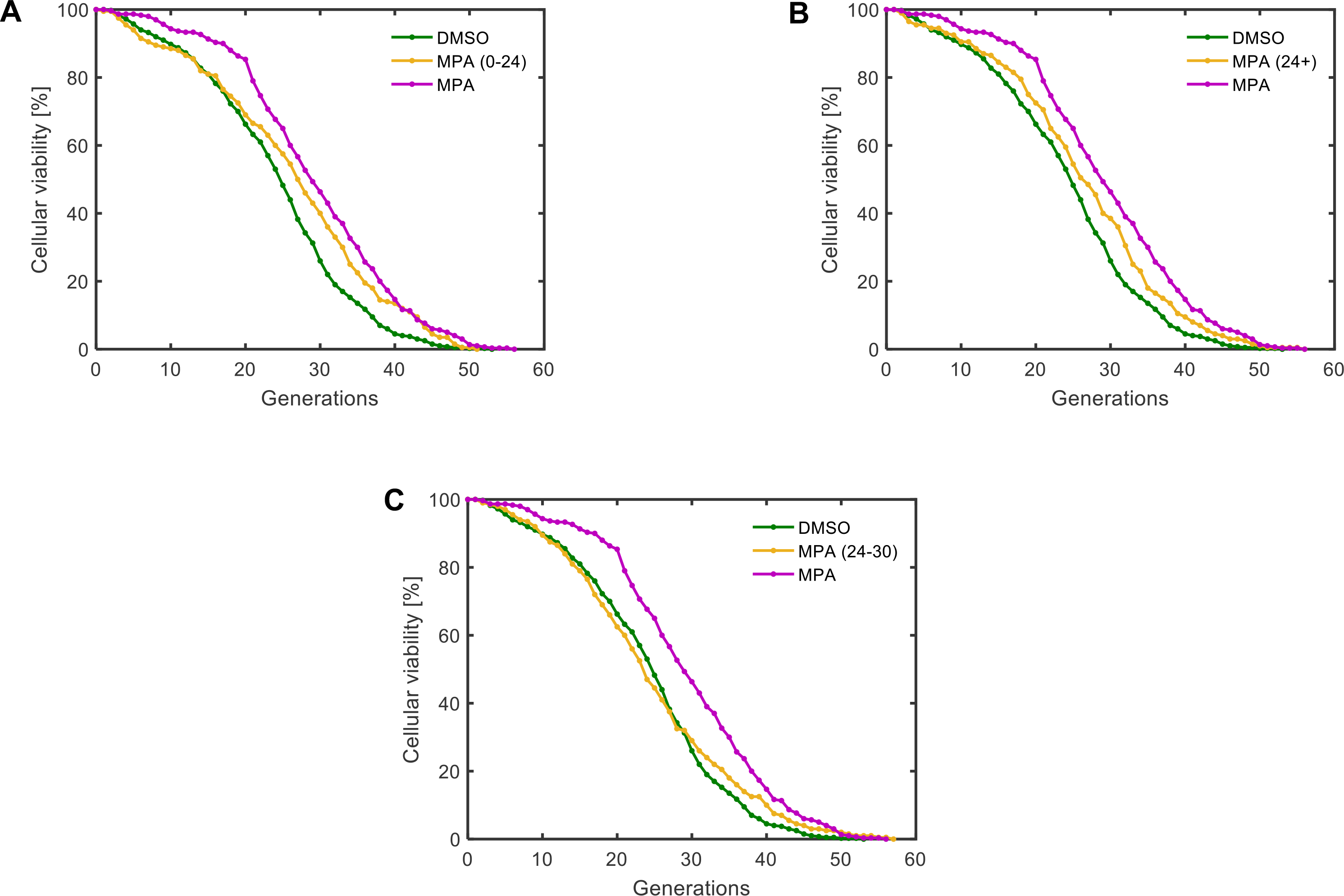
MPA slows, rather than reverses, the aging process. **A-C.** Replicative lifespan curves for *S. cerevisiae* treated with 10 μM MPA only during the first 24 hours of a Replicator experiment (**A**), only after the first 24 hours of a Replicator experiment (**B**), or between the 24^th^ and 30^th^ hour of a Replicator experiment (**C**). N = 400 cells for the DMSO condition, pooled from four independent experiments of 100 cells each; N = 300 cells for the full-time MPA condition, pooled from three independent experiments of 100 cells each; N = 200 cells for the partial-time MPA conditions, pooled from two independent experiments of 100 cells each.

### Assessing cellular processes in which GMP and its metabolites could regulate lifespan

As part of further characterizations to understand the means by which GMP synthesis inhibition extends lifespan, we considered the cellular processes in which GMP and its downstream metabolites are involved. Among the many downstream metabolites, guanine derivatives are used for transcription, signal transduction, and as an energy source for enzymatic reactions.

While GTP is used as an energy source for some enzymatic reactions and also during some signal transduction processes involving G proteins, these uses do not consume the nucleoside, only the high-energy phosphate bonds, and the resulting GDP or GMP is readily recycled by phosphotransferases inside the cell back into GTP. Thus, we do not expect signal transduction or energy source hypotheses to contribute much to the observed lifespan extension, especially since the dosage of MPA applied did not substantially impede cell growth.

To see if cell-to-cell differences in lifespan could be explained by potential differences in transcriptional activity, we analyzed single-cell microscopy data in two different wild-type strains carrying two copies of a fluorescent reporter driven by the constitutive *TEF1* promoter integrated at the *HIS3* locus. The only difference between the two strains is that the first strain uses a stable EGFP reporter, while the second strain uses semistable GFP (ssGFP), which is a modified EGFP with a degron from the cyclin Cln2 attached^22^. For each strain, the replicative lifespans of 50 cells were measured, and the average expression level of the reporter for each cell were recorded with hourly snapshots; for each cell, the fluorescence value recorded in the snapshots were averaged to produce the average expression level throughout its replicative lifespan. We found no correlation between average fluorescence and RLS on a single-cell level in the EGFP strain (Fig. 5A; Pearson’s *r* = −0.0051). On the other hand, the ssGFP strain displayed significant negative correlation between the average fluorescence level and RLS (Fig. 5B; Pearson’s r = −0.3557; Holm-Bonferroni adjusted *p* = 0.0337). To test the generality of the observed negative correlation in the ssGFP strain, we performed the same analysis in three additional strains that changed the locus of integration (Fig. 5C), the number of reporter copies (Fig. 5D), and the promoter used to drive the reporter (Fig. 5E). We found the same negative correlation in all three strains (Pearson’s *r* ranging from −0.3387 to −0.4866; Holm-Bonferroni adjusted p-values < 0.05).

**Figure 5.**
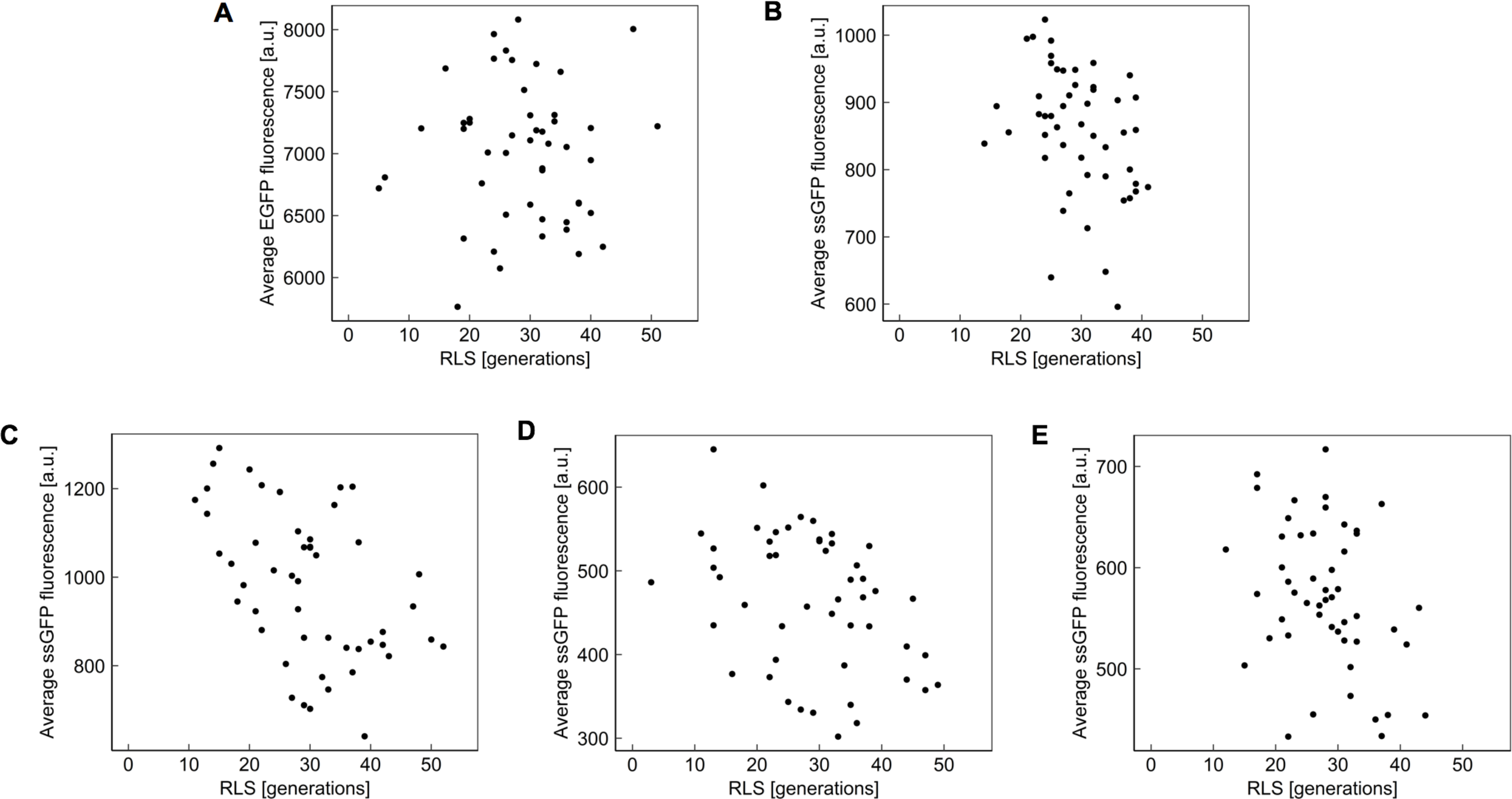
Correlation analysis between average single-cell expression level and replicative lifespan of cells for five wild-type diploid strains. **A.** Homozygous P_*TEF1*_-EGFP reporter at *HIS3* locus (Pearson’s *r* = −0.0051; Holm-Bonferroni adjusted *p* = 0.9718). **B.** Homozygous P_*TEF1*_-ssGFP reporter at *HIS3* locus (Pearson’s *r* = −0.3557; Holm-Bonferroni adjusted *p* = 0.0337). **C.** Homozygous P_*TEF1*_-ssGFP reporter at *SAM2* locus (Pearson’s r = −0.4866; Holm-Bonferroni adjusted *p* = 0.0017). **D.** Heterozygous P_*TEF1*_-ssGFP reporter at *SAM2* locus (Pearson’s *r* = −0.3765; Holm-Bonferroni adjusted *p* = 0.0282). **E.** Homozygous P_*PGK1*_-ssGFP reporter at *HIS3* locus (Pearson’s *r* = −0.3387; Holm-Bonferroni adjusted *p* = 0.0337).

The concentration of a protein in an aging cell is affected by the transcription, translation, and degradation rates of the protein, as well as by any age-associated global changes on these rates. Since EGFP is a stable protein, its degradation is virtually non-existent. Thus, its concentration is a faithful reporter of the historical average transcription and translation levels in the cell. The absence of any correlation between EGFP level and RLS (Fig. 5A) thus indicates that the global level of transcription and translation together has no effect on the cell’s replicative lifespan.

The four strains expressing ssGFP, on the other hand, do display a negative correlation between ssGFP levels and lifespan at the single-cell level. Since global transcription and translation levels together do not measurably impact lifespan (Fig. 5A), it follows that difference in degradation of ssGFP is likely responsible for the correlation observed. Further supporting this view, the truncated Cln2 degron attached to ssGFP targets it for degradation via the proteasome^23^, and it has previously been shown that activation of the proteasome pathway increases RLS^2,3^. Thus, the observed negative correlation is expected to be due to the fact that cells with higher levels of proteasome activity (which increase lifespan) also degrade ssGFP more rapidly, resulting in lower levels of ssGFP fluoroscence. Taken together, based on data collected from longitudinally-tracked aging cells, these results provide single-cell level support on the conclusion that cells with higher proteasome activity live longer.

## DISCUSSION

In this study, we showed how MPA, an FDA-approved immunosuppressant, facilitates lifespan extension in yeast. Since MPA has a well-characterized role on GMP metabolism, a direct mechanism for its lifespan extension effect emerged through GMP synthesis inhibition. This was further confirmed by the observation that adding excess guanine into growth media reverses the RLS extension effect of MPA. Here we also showed that activated proteasome extends lifespan in part through GMP insufficiency. Taken together, our work elucidates the involvement of nucleotide metabolism in the aging process.

MPA exerts its lifespan-extending effect by slowing, rather than reversing, the aging process. As part of our characterizations to locate the pathway(s) through which MPA impacts lifespan, we find that the nutrient-sensing and sirtuin pathways are not involved in lifespan extension from the use of MPA. Evaluating a set of cellular processes or phenotypes for their potential role in MPA-caused lifespan extension, we conclude that the lifespan extension effect of MPA is not facilitated through age-associated global changes in gene expression (transcription/translation combined).

Throughout this study, we used MPA at the 10 μM concentration because of the “standard” nature of this concentration based on many compound-screening assays. However, a dose-response characterization we performed showed that 10 μM was indeed the most efficient MPA concentration in terms of its RLS-extending effect (Fig. S7).

Our work uncovers the possibility of a new class of therapeutics targeting aging-related disease. Future studies will determine the clinical viability of GMP synthesis inhibitors for delaying the onset of or for the prevention of aging-related disease. The finding that mycophenolic acid extends lifespan in a *ΔTOR1* genetic background promotes the idea of combination therapy with the TOR inhibitors currently in clinical trials. Characterization of new targets through which cellular aging^20,21,22,24^ can be regulated represents an important milestone towards the ultimate goal of fully understanding aging and lifespan determinants at single-cell resolution.

## MATERIALS and METHODS

### Yeast strains and media conditions used for determining haploid lifespan characteristics, proteasome activity, and GFP expression

All experiments in haploid *S. cerevisiae* were conducted in a BY4741 strain background (TransOMIC TKY0002). Haploid deletion strains were purchased from the Yeast Knockout Collection maintained by GE Dharmacon. The haploid Δ*SIR2*Δ*FOB1* strain was constructed by deletion of *FOB1* from a Δ*SIR2* background using the lithium acetate method^25^ and a *HIS5* selectable marker. The haploid SIR2-OE strain was prepared by integration of the native *SIR2* promoter, gene, and terminator at the *URA3* locus under URA selection. The haploid strain carrying P_*TEF1*_-GFP was constructed by integration of the *TEF1* promoter, eGFP, and terminator at the *HIS3* locus under URA selection. Complete supplement mixture (CSM 2% glucose) media was used for all experiments, with cells maintained in aerobic conditions at 30 °C in 50 mL conical tubes (Becton Dickinson F2070) or sterile 250 mL Erlenmeyer flasks. Cultures were performed in an Innova-42 shaker (New Brunswick Scientific) set to 225 rpm.

### Replicative lifespan measurements for haploid yeast in the Replicator

RLS measurements were performed in the Replicator device for haploids, as described in our previous publication^11^. Briefly, cells were grown overnight in CSM 2% glucose for 24 hours to an OD_600_ of 0.1, and then loaded to the microfluidic Replicator device. Media was then flowed through the device, containing DMSO (American Bio AB00435), 10 μM mycophenolic acid (Sigma M5255), and/or guanine (Sigma G11950) at a saturating concentration of 20 mg/L. An automated microscope was used to collect time-lapse images of individual mother cells every 10 min for 120 hours, and RLS was determined by counting the number of daughters produced before death. RLS was scored manually by observing the time-lapse image series produced in a Replicator experiment. Cells were included for measurement if they entered the trap prior to the 10^th^ hour of the experiment. Passage into each generation was denoted by the appearance of a bud.

### Protein extraction and proteasome assay

For protein extraction, 50 mL of cells were grown for 18 hours to an OD600 of approximately 0.8, then transferred to a 50 mL conical tube and centrifuged at 4255 xg for 5 minutes at room temperature. The supernatant was discarded, and the cells were resuspended in 150 μL of cold lysis buffer (50 mM Tris-HCl, pH 7.5, 0.5 mM EDTA, 5 mM MgCl_2_, with cOmplete™ ULTRA mini protease inhibitor tablets, EDTA free) and transferred to a 1.5 mL tube. A ¼ volume of 500-750 μm glass beads (Acros Organics 397641000) was added to each tube. For 10 rounds, the tubes were chilled in ice water for 1 minute, then vortexed at maximum speed for 30 seconds to physically rupture the cells, and returned to the ice water. Samples were then spun for 3 minutes at 2500 xg and 4 °C, and the supernatant was transferred to a fresh tube. The solution was further clarified by centrifugation at 8000 xg for 10 minutes at 4 °C, and the supernatant was transferred to a fresh tube. Protein concentration was measured using a Nanodrop measuring the absorbance of the sample at 280 nm.

The proteasome assay used was previously described^19^. The assay was performed in a 96-well clear-bottom plate (Costar 3603) with 50 μg of total protein in 200 μL of lysis buffer. The fluorogenic proteasome substrate Suc-LLVY-AMC (Bachem I-1395) was added to a final concentration of 100 μM. Fluorescence intensity with an excitation wavelength of 380 nm and an emission wavelength of 460 nm was recorded at 5-minute intervals using a Neo2 plate reader (BioTek) set to mix constantly and maintain 30 °C. Negative control reactions were performed in the presence of 50 μM MG132 (Sigma 474787), a proteasome inhibitor.

### Total RNA measurements

In four different growth conditions (containing DMSO (American Bio AB00435), DMSO plus 10 μM mycophenolic acid (Sigma M5255), DMSO plus guanine (Sigma G11950) at a saturating concentration of 20 mg/L, DMSO plus 20 mg/L guanine plus 10 μM mycophenolic acid) in CSM minimal media (2% glucose), haploid yeast cells of BY4741 genetic background were grown (in triplicates) for 18 hours in 10 mL volume to an OD_600_ of 1.0 ± 0.14. Cells were then spun at 1500g at 4°C for 3 minutes. Pellets were resuspended in 1 mL of ice-cold water and spun again at 4°C. Pellets were then resuspended in 400 μl of TES solution, and 400 μl of acid phenol (pH 4.9) were added. Cells were vortexed and incubated at 65 °C for 1 hour with occasional vortexing. Lysates were then placed on ice for 5 minutes followed by a 5 minute spin at 4°C. The aqueous phase was transferred to a clean 1.5 mL tube, and 400 μl acid phenol was added. Tubes were then vortexed and spun for 5 minutes at 4°C. This step was carried out twice. The aqueous phase was then again transferred to a 1.5 mL tube and 400 μl of chloroform were added. Samples were vortexed and spun for 5 minutes at 4°C. The aqueous phase was transferred to a new 1.5 mL tube and 40 μl of 3 M sodium acetate (pH 5.3) and 1 mL of ice-cold 100% ethanol were added. Samples were again spun for 5 minutes at 4°C and the RNA pellet was washed with ice-cold 70% ethanol. Pellet was resuspended in 25 μl of nuclease free water and stored overnight at −20°C. RNA was then quantified using a NanoDrop™ 2000 Spectrophotometer (ThermoScientific). RNA concentration of each sample was normalized to each sample’s OD_600_ density measured right after the 18 hours growth period.

### Growth conditions and flow cytometry for GFP measurements

All GFP measurements from the wild type BY4741 and the P_*TEF1*_-GFP-containing strain were performed using a BD FACSVerse, with cells grown, in triplicates, for 18 hours to mid-log phase, an OD_600_ of 0.1-0.5, at the start of the FACS measurement. Cultures were placed on ice after 18 hours, then immediately used for flow cytometry. Complete supplement mixture (CSM 2% glucose) media was used for all experiments. Media conditions during the 18 hours growth also included: 1. DMSO (American Bio AB00435); 2. DMSO and 10 μM mycophenolic acid (Sigma M5255); 3. DMSO and guanine (Sigma G11950) at a saturating concentration of 20 mg/L; 4. DMSO, 10 μM mycophenolic acid, and 20 mg/L guanine. Cells were maintained in aerobic conditions at 30 °C in 50 mL conical tubes (Becton Dickinson F2070). Cultures were performed in an Innova-42 shaker (New Brunswick Scientific) set to 225 rpm.

### Experiments performed using diploid yeast cells

To measure single-cell level correlations between average fluorescence intensity and replicative lifespan, we reanalyzed data generated for a previous paper^22^. The description and construction of the diploid yeast strains, media conditions, set-up of the aging experiments, and data analysis steps are described in detail in that paper^22^. The five diploid strains from which we collected single-cell average fluorescence data across each cell’s lifespan were: strain heterozygous in P_*TEF1*_-ssGFP cloned at *SAM2* locus, strain homozygous in P_*TEF1*_-ssGFP cloned at *SAM2* locus, strain homozygous in P_*TEF1*_-ssGFP cloned at *HIS3* locus, strain homozygous in P_*TEF1*_-eGFP cloned at *HIS3* locus, and strain homozygous in P_*PGK1*_-ssGFP cloned at *HIS3* locus. ssGFP indicates semi-stable GFP^22^.

### Statistical methods for lifespan characteristics, proteasome activity, GFP expression, RNA levels

Differences in lifespan characteristics were assessed through Log-Rank test using MATLAB with a cut-off value of P = 0.05. A summary of the statistical tests and their results is provided in Table S1. The script for Log-Rank test was downloaded from MATLAB central: http://www.mathworks.com/matlabcentral/fileexchange/20388-logrank. Differences in proteasome activity, P_*TEF1*_-GFP expression, and RNA levels were assessed using the unpaired, two-tailed, parametric t-test function.

## ACKNOWLEDGEMENTS

We thank Shirin Bahmanyar, Marc Hammarlund, Gerald Shadel, Patrick Sung, and Acar Lab members for comments and feedback on different stages of this work. EAS acknowledges support through an NSF Graduate Research Fellowship and Gruber Science Fellowship. GU acknowledges support through an NIH pre-doctoral training grant and Gruber Science Fellowship. MA acknowledges funding from the Ellison Medical Foundation (AG-NS-1015-13) and US National Institutes of Health (1DP2AG050461-01 and 1R01GM127870-01).

## AUTHOR CONTRIBUTIONS

EAS, PL, and MA contributed to project planning, and design and preparation of the manuscript. EAS and PL contributed to strain construction, data collection, data analysis, and preparation of the figures. GU performed the total RNA quantification experiments and collected their data. TTO contributed to strain construction, data collection, and data analysis. TZY contributed to data analysis and writing. RS performed the correlation analysis for the data collected from diploid yeast, plotted the results, and contributed to writing. EAS, PL, GU, TTO, TZY, RS, and MA interpreted the data and results, and read and approved the manuscript.

