## Supplementary Information for "Inhibition of GMP synthesis extends yeast replicative lifespan"

**I. Supplementary Text**

**II. Supplementary Figures S1-S7**

**III. Supplementary Tables S1-S2**

**IV. Supplementary References**

### I. SUPPLEMENTARY TEXT

#### Longevity Placement Test

Longevity Placement Test (LPT) is a mutually exclusive, collectively exhaustive method to determine the relationship of a longevity intervention to a known genetic regulator of lifespan. For a given intervention that extends lifespan, any of three possible relationships may exist relative to a known genetic regulator of lifespan: (1) the intervention may act to extend lifespan independently from the known regulator; (2) the intervention may act downstream from the known regulator, converging on a single component of the known regulator's lifespan pathway; (3) the intervention may act upstream from or upon the known regulator, ultimately modulating lifespan through the genetic regulator.

In Step 1 of the LPT, the longevity intervention is applied to a strain in which a genetic regulator of lifespan is deleted. In the event that lifespan extension from the longevity intervention is observed in this background, only possibilities (1) and (2) above remain valid (Fig. S1A). If no lifespan extension is observed, then possibilities (2) and (3) remain valid. Possibility (2) cannot be ruled out in this step, since non-saturating action by the genetic regulator could leave room for lifespan extension by the longevity intervention, while saturating action would preclude it.

In Step 2 of the LPT, an epistatic agent, which prevents lifespan extension from the longevity agent, is applied to a strain in which some upstream member of the genetic regulator's lifespan pathway has been modified to extend lifespan. This step differentiates possibility (2) from the remaining possibility after Step 1. In the event that lifespan extension from the genetic regulator's pathway is suppressed by the epistatic agent, this determines that possibility (2) is correct. In the event that no epistasis is observed, the remaining possibility, (1) or (3), is correct (Fig. S1B).

Preconditions, both physical and experimental, exist for an LPT experiment to conclusively relate a longevity intervention to a given pathway. Two physical factors must exist for an exhaustive LPT test: a longevity agent to test, and an epistatic agent that prevents lifespan extension from the longevity agent. Importantly, the epistatic agent must not directly affect the longevity agent, such as through inactivating it. Ideally, it should exert an opposing effect on some downstream target, ensuring that its suppression will be generalizable to any actor upstream of the longevity agent's target. There also exist constraints on the choice of genetic manipulations for investigation. For Step 1, probe gene deletion should not shorten RLS, as this may mask longevity effects[1]. For Step 2, the intervention which extends RLS must act upon or upstream from the Step 1 probe gene in order to create complete coverage of the pathway. However, the intervention in Step 2 need not always be a gene deletion; for example, in the case of SIR2, deletion of SIR2 in Step 1 could be complemented by overexpression of SIR2, an intervention which extends lifespan, in Step 2.

### II. SUPPLEMENTARY FIGURES

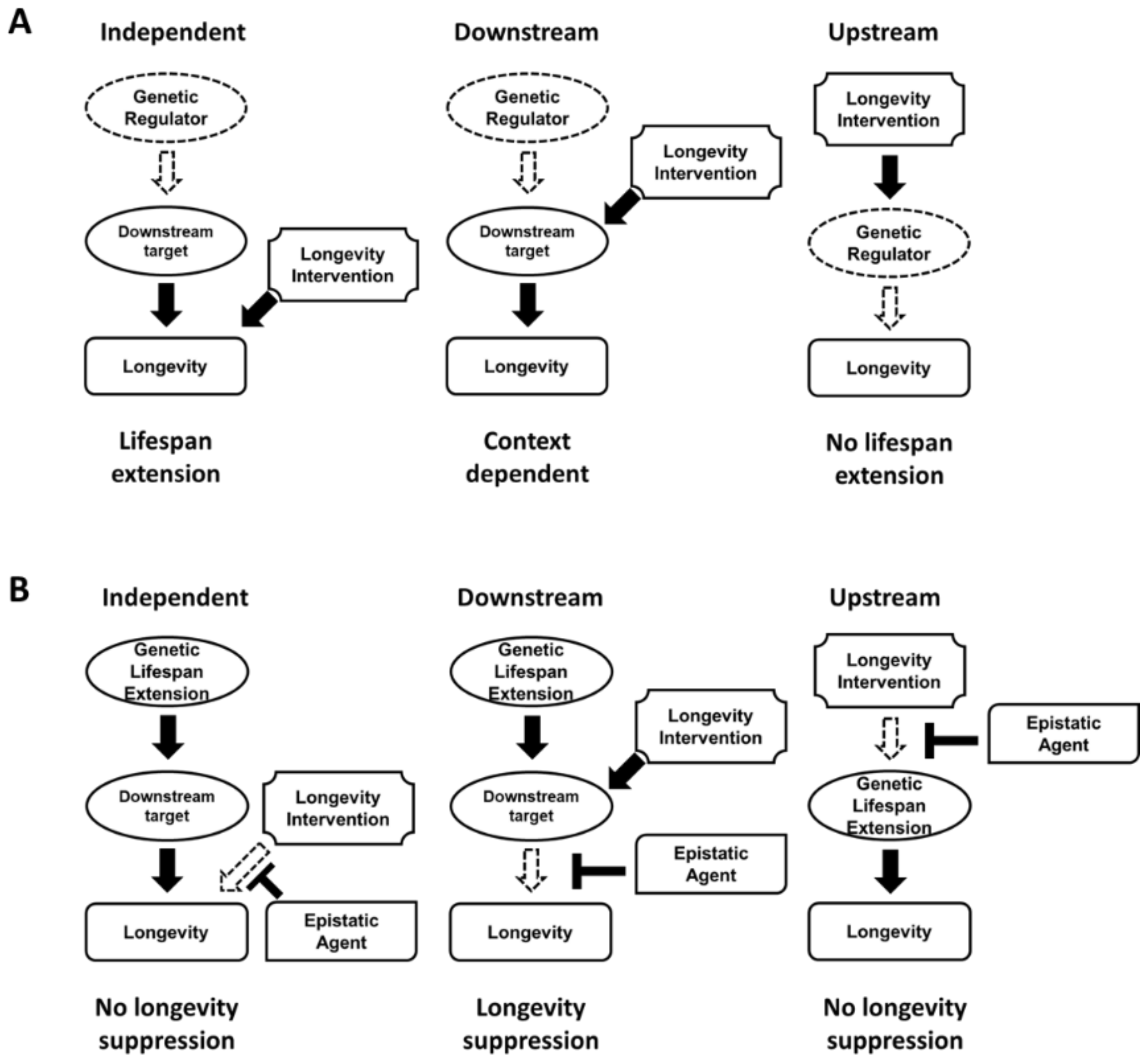


**Figure S1. Possible outcomes of the longevity placement test.** Possible network architectures, and the outcomes of the LPT in Step 1 (**A**) and Step 2 (**B**). Interactions shown in dashed lines represent those that are prevented, either by the suppression agent, or via deletion of the gene.

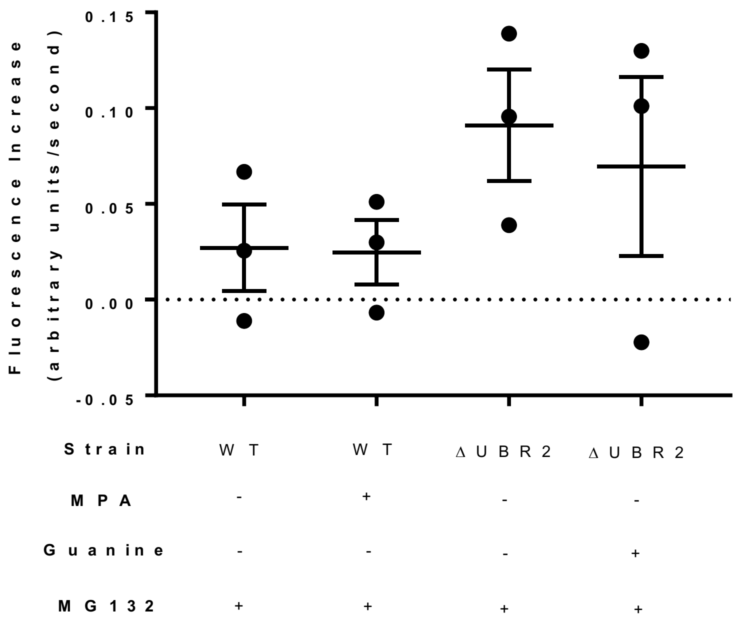


**Figure S2. Negative control for proteasome activity measurement.** The proteasome activity experiment shown in Fig. 3C is controlled for by adding MG-132, a proteasome inhibitor, to separate wells of the experiment run concurrently. The low rate of fluorescence increase in the presence of MG-132 indicates that our measurements were specific to the proteasome.

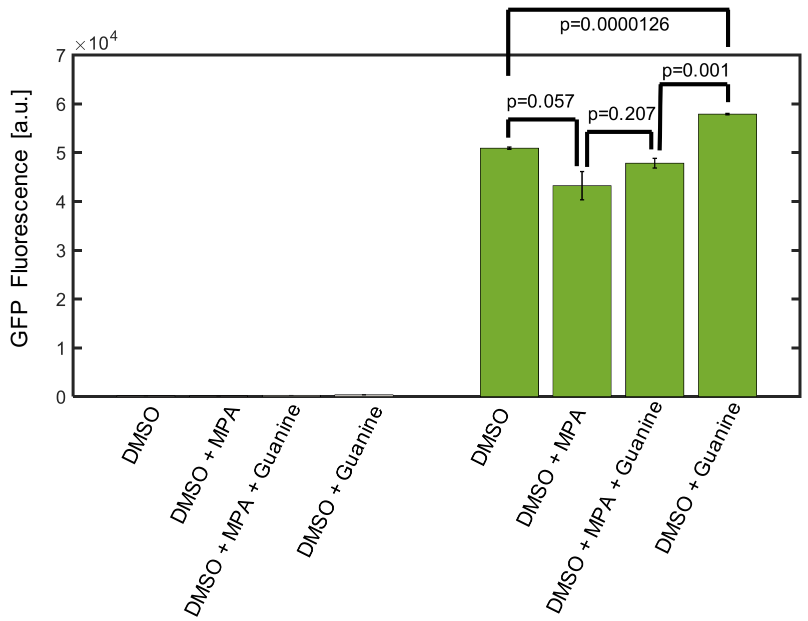


**Figure S3. Flow cytometry-measured mean GFP levels.** Haploid yeast cells were grown in complete supplement mixture (CSM 2% glucose) supplemented with the media components as shown in the panel. The four data columns shown on left were obtained from the wild-type strain (BY4741 with no reporter), while the four data columns on right were obtained from the strain carrying $P_{TEF1}$-GFP reporter in a BY4741 genetic background. Error bars denote SEM (N=3).

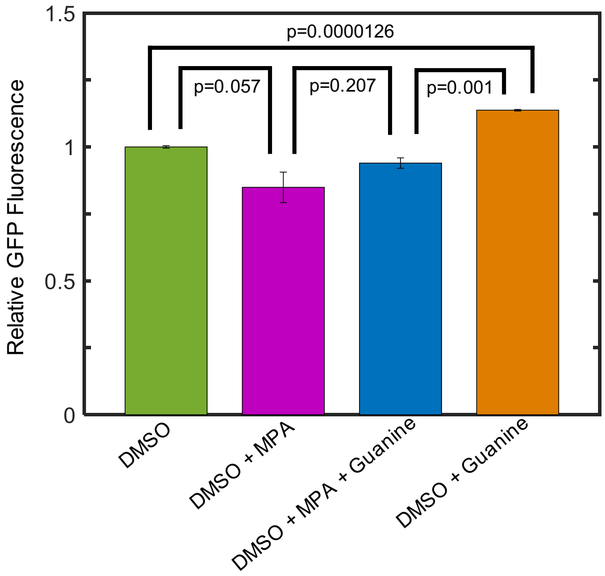


**Figure S4. MPA reduces gene expression levels.** Flow cytometry-measured GFP levels relative to the mean GFP level of the DMSO environment. Haploid yeast cells were grown in complete supplement mixture (CSM 2% glucose) supplemented with the media components as shown in the panel. The data were obtained from the strain carrying $P_{TEF1}$-GFP reporter in a BY4741 genetic background. Error bars denote SEM (N=3).

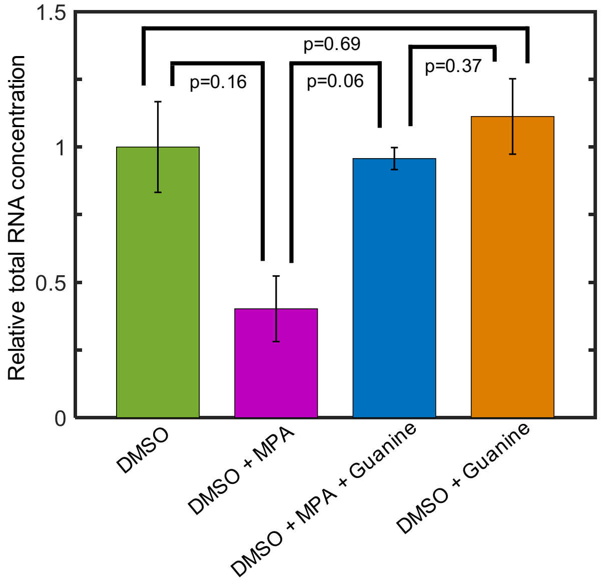


**Figure S5. MPA reduces relative total RNA levels.** Total RNA levels relative to the DMSO control. Haploid wild-type yeast cells (BY4741) were grown in complete supplement mixture (CSM 2% glucose) supplemented with the media components as shown in the panel. For each growth condition/environment, total RNA levels were measured from three independently grown cultures. The total RNA level of each independent culture was normalized by the mean total RNA level obtained in the DMSO environment, followed by calculating the mean and SEM (N=3) of the normalized RNA levels for each environment. RNA concentration of each sample was normalized to each sample's $OD_{600}$ density measured right after the 18 hours growth period.

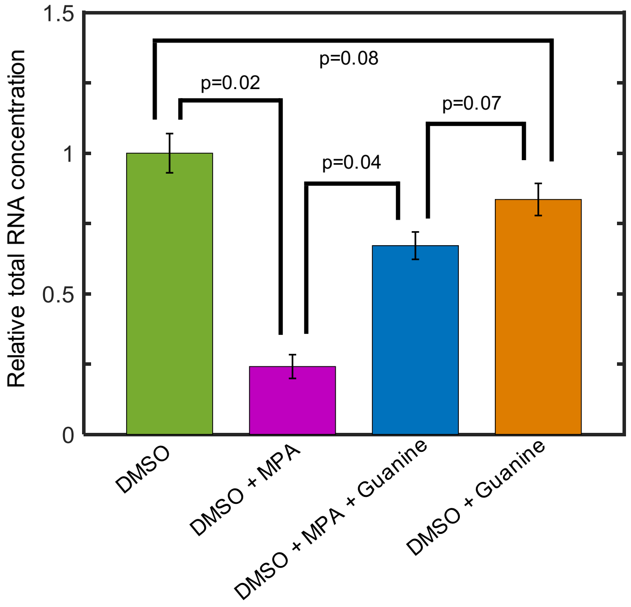


**Figure S6. Relative total RNA levels measured in the *UBR2*Δ cells.** Total RNA levels relative to the DMSO control. Haploid *UBR2*Δ yeast cells were grown in complete supplement mixture (CSM 2%

glucose) supplemented with the media components as shown in the panel. For each growth condition/environment, total RNA levels were measured from three independently grown cultures. The total RNA level of each independent culture was normalized by the mean total RNA level obtained in the DMSO environment, followed by calculating the mean and SEM (N=3) of the normalized RNA levels for each environment. RNA concentration of each sample was normalized to each sample's $OD_{600}$ density measured right after the 18 hours growth period.

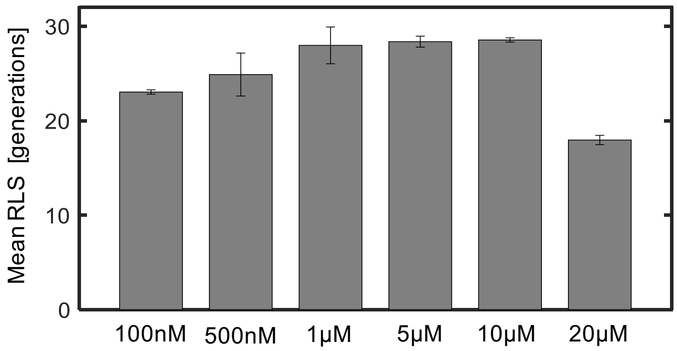


**Figure S7. Dose-response characterization of MPA.** Mean replicative lifespan across multiple MPA concentrations. Error bars denote S.E.M. (N=2-3). For each MPA concentration, mean RLS from 2 biological replicates was calculated, except for 10μM MPA, for which mean RLS from 3 biological replicates was calculated. Each replicate contributed 100 cells, except for 500nM MPA, for which there were 95 and 105 cells contributed by the two replicates.

### III. SUPPLEMENTARY TABLES

| Condition 1 | Condition 1 Mean Lifespan (Generations) | Condition 2 | Condition 2 Mean Lifespan (Generations) | Percent Change | p-value obtained from log-rank test |
|---|---|---|---|---|---|
| BY4741-DMSO | 23.39 | BY4741-Guanine | 22.7 | -2.94997862 | 0.9034 |
| BY4741-DMSO | 23.39 | BY4741-MPA | 28.553 | 22.0735357 | 1.43E-09 |
| BY4741-DMSO | 23.39 | BY4741-MPA-Guanine | 24.183 | 3.390337751 | 0.0463 |
| BY4741-DMSO | 23.39 | ΔHXK2-DMSO | 26.845 | 14.77126977 | 6.60E-04 |
| ΔHXK2 -DMSO | 26.845 | ΔHXK2 -Guanine | 27.6 | 2.812441795 | 0.45 |
| ΔHXK2-DMSO | 26.845 | ΔHXK2-MPA | 31.35 | 16.78152356 | 0.0000269 |
| BY4741-MPA | 28.553 | ΔHXK2-MPA | 31.35 | 9.795818303 | 0.0039 |
| BY4741-DMSO | 23.39 | ΔTOR1-DMSO | 28.805 | 23.1509192 | 4.27E-08 |
| ΔTOR1 -DMSO | 28.805 | ΔTOR1 -Guanine | 28.935 | 0.451310536 | 0.87 |
| ΔTOR1-DMSO | 28.805 | ΔTOR1-MPA | 32.525 | 12.91442458 | 0.000634 |
| BY4741-MPA | 28.553 | ΔTOR1-MPA | 32.525 | 13.91097258 | 2.40E-06 |
| BY4741-DMSO | 23.39 | SIR2O.E.-DMSO | 28.595 | 22.25309962 | 1.09E-09 |
| SIR2O.E.-DMSO | 28.595 | SIR2O.E.-Guanine | 27.31 | -4.49379262 | 0.25 |
| ΔSIR2ΔFOB1-DMSO | 25.025 | ΔSIR2ΔFOB1-MPA | 27.125 | 8.391608392 | 0.0123 |
| BY4741-MPA | 28.553 | ΔSIR2ΔFOB1-MPA | 27.125 | -5.00122579 | 0.762 |
| BY4741-DMSO | 23.39 | ΔUBR2-DMSO | 28.8 | 23.12954254 | 2.32E-13 |
| ΔUBR2 -DMSO | 28.8 | ΔUBR2 -Guanine | 27.28 | -5.27777778 | 0.0147 |
| ΔUBR2-DMSO | 28.8 | ΔUBR2-MPA | 31.5 | 9.375 | 0.0122 |
| BY4741-MPA | 28.553 | ΔUBR2-MPA | 31.5 | 10.32115715 | 2.35E-04 |
| BY4741-Guanine | 22.7 | ΔUBR2 -Guanine | 27.28 | 20.17621145 | 7.99E-08 |
| BY4741-DMSO | 23.39 | ΔPRE9-DMSO | 23.005 | -1.64600257 | 0.5019 |
| ΔPRE9-DMSO | 23.005 | ΔPRE9-MPA | 27.84 | 21.01717018 | 0.0000684 |
| BY4741-MPA | 28.553 | ΔPRE9-MPA | 27.84 | -2.49711064 | 0.1524 |
| BY4741-DMSO | 23.39 | BY4741-MPA(0-24) | 24.37 | 4.189824711 | 0.0013 |
| BY4741-DMSO | 23.39 | BY4741-MPA(24+) | 25.55 | 9.23471569 | 0.0066 |
| BY4741-DMSO | 23.39 | BY4741-MPA(24-30) | 23.755 | 1.560495938 | 0.2429 |
| BY4741-MPA | 28.553 | BY4741-MPA(0-24) | 24.37 | -14.6499492 | 0.0436 |
| BY4741-MPA | 28.553 | BY4741-MPA(24+) | 25.55 | -10.5172836 | 0.0122 |
| BY4741-MPA | 28.553 | BY4741-MPA(24-30) | 23.755 | -16.8038385 | 4.83E-04 |
| BY4741-MPA(0-24) | 24.37 | BY4741-MPA(24-30) | 23.755 | -2.52359458 | 0.15 |
| BY4741-MPA(0-24) | 24.37 | BY4741-MPA(24+) | 25.55 | 4.842018876 | 0.66 |
| BY4741-MPA(24+) | 25.55 | BY4741-MPA(24-30) | 23.755 | -7.02544031 | 0.279 |

**Table S1. Summary of lifespans and log-rank tests.**

| Genotype | Marker |
|---|---|
| ΔSIR2/ΔFOB1 | KanMX/HIS5 |
| SIR2-OE | URA3 |
| ΔHXK2 | KanMX |
| ΔTOR1 | KanMX |
| ΔUBR2 | KanMX |
| ΔPRE9 | KanMX |
| $P_{TEF1}$-GFP | URA3 |

**Table S2. Markers used in strains.** The *S. cerevisiae* strains used in this study have the BY4741 genetic background. Markers used for gene deletion or insertion in haploid yeast (as described in the Materials and Methods section) are shown in the above table.
